# Clinical and Genetic insights in a tertiary care center cohort of patients with bicuspid aortic valve

**DOI:** 10.1101/2021.02.15.431243

**Authors:** Alexis Théron, Anissa Touil, Noémie Résseguier, Gwenaelle Collod-Beroud, Giulia Norscini, Anne-Sophie Simoni, Gaёlle Odelin, Frédéric Collart, Stéphane Zaffran, Jean-François Avierinos

**Author notes:** **Co-corresponding authors:** Jean-François Avierinos, Stéphane Zaffran.

## Abstract

**BACKGROUND:** to describe spectrum of valve function in bicuspid aortic valve (BAV) patients referred to a tertiary care center and to investigate genetic pathways associated with valve degeneration.

**METHODS:** All consecutive patients with BAV were prospectively included from 2014 to 2018. BAV was defined according to embryologic and Sievers classifications. Clinical and echo variables were compared according to aortic valve function. Aortic valve tissues were collected from BAV patients (n=15) operated for severe aortic stenosis (AS-BAV, n=7) or aortic regurgitation (AR-BAV, n=8). RT-qPCR was performed to compare gene expression level in AS-BAV, AR-BAV and controls corresponding to healthy tricuspid aortic valves collected on human heart explant immediately after transplantation (n=5).

**RESULTS:** Out of 223 adults with BAV, mean age 53±17 years, 83% had left-right coronary cusps fusion, 80% Sievers type 1 BAV and 49% aortopathy. Twenty-four patients had normal valve function, 66 patients had AS-BAV, 91 patients had AR-BAV and 40 patient had AR+AS BAV. BAV phenotype did not predict neither AS-BAV nor AR-BAV (all p>0.1). By multivariable analysis, age >50 (41.6[10.3-248.2],p<0.001) and presence of raphe(12.8[2.4-87.4],p<0.001) were significantly associated with AS-BAV and male gender of AR-BAV(5[1.6-16.4], p=0.005). RT-qPCR revealed overexpression of *RUNX2* in AS-BAV (17.67±1.83 vs 3.25±0.93, p=0.04), and overexpression of *COL1A1* (4.01±0.6vs2.25±0.5,p=0.03) and *FLNA* (23.31±7.5vs1.97±0.3,p=0.03) in AR-BAV.

**CONCLUSIONS:** This prospective study confirmed high prevalence of valve dysfunction at first diagnosis of BAV in a referred population. Clinical and echo variables are poorly associated with BAV dysfunction. The leaking or stenotic processes could be both supported by dysregulation of specific genetic pathways

## INTRODUCTION

Congenital bicuspid aortic valve(BAV) is defined as a spectrum of altered aortic valves presenting with two functional cusps and less than three zones of parallel apposition without fusion.^1^ It is the most frequent congenital heart disease (CHD) affecting 0.5 to 2% of the population with a strong male predominance.^2^ BAV is associated with excess mortality in some cases^3^ and accounts for more complications than all other CHDs combined,^4,5 6^ related to increased risk of ascending aorta aneurism, dissection and valve degeneration, the latter occurring 15 to 20 years earlier as compared to tricuspid aortic valve.^7,8^ Aortic valve and/or ascending aorta replacements are the only therapeutic option,^9,10^ with 25% incidence of this end-point at 20 years in patients without significant valvular dysfunction or aorta dilatation at first diagnosis.^7^ Predictors of surgery include age>50 years and presence of valve degeneration at diagnosis, associated with a 4-fold increased risk of valve replacement at 20 years.^7^ However, mechanisms underlying degeneration from normal BAV to leaking or stenotic BAV remain poorly understood.^2 11 12^

The Bicuspid Aortic valve Project (BAP) is designed to collect clinical and genetic data on consecutive patients prospectively admitted with a definite diagnosis of BAV. We sought to describe the clinical and echocardiographic spectrum of consecutive BAV patients as they present in a tertiary care center and to determine factors associated with each type of valve degeneration.

## METHODS

### Study population

The BAP is a French single centre prospective study including consecutive patients with non-syndromic BAV for clinical and genetic purpose. Patients were recruited between 2014 and 2018 in La Timone Hospital, Marseille, France. Patients were not included if they had connective tissue disease, previous aortic valve surgery and less than 18 years old. This study was performed in accordance with institutional guidelines. Written consent was obtained from all patients.

### BAV diagnosis

Definite diagnosis of BAV was based on visualization in short axis view of two functional cusps in mid-systole; partial fusion between cusps was considered as BAV. TEE was performed when diagnosis was uncertain after TTE. Raphe was considered as definitely present when clearly visible by TTE or TEE. BAV phenotypes were specifically classified regarding both Siever’s classification and embryologic classification^1,6^ and all echocardiograms were performed by two experienced cardiologists (JFA,AT) to ensure homogeneous collection of data.

### BAV classification

Sievers classification is based on presence or absence of raphe between fused cusps. Embryologic classification refers to type of embryologic cusp fusion -or non-separation-based on commissural axis orientation in short axis view. Typical BAV is defined by 11-17 or 10-16 o’clock positions of commissural axis, corresponding to left-right coronary cusps fusion (L-R BAV). Atypical BAV is defined by 13-19 or 12-18 o’clock positions of commissural axis, corresponding to non-coronary and right coronary cusps fusion (NC-R). The rare non coronary-left coronary cusps fusion (NC-L) is also labeled atypical and defined by 14-20 o’clock position

### Baseline Clinical data

Clinical data were prospectively collected at the time of inclusion as well as family history of cardiac surgery, aortic dissection and CHD. Surgical correction of valve dysfunction and associated aortopathy was indicated according to the European and US guidelines.^10,9^

### Baseline Echocardiographic data

Aortic regurgitation (AR) was graded through an integrative approach.^10,9^ Mechanism of AR was separated into prolapse of the fused leaflet, cusps restriction or both. Aortic stenosis (AS) was considered as present when mean transvalvular gradient>10mmHg with a restriction of the aortic valve opening. Severe AS was defined by a mean gradient>40mmHg and an indexed AVA<0,6cm^2^/m^2^.^9^ Normal valve function was defined by the absence of AR even mild and of AS with a mean trans-valvular gradient<10mmHg. Associated aortopathy was defined by an ascending aorta diameter≥40mm and separated into “root phenotype”, “tubular phenotype” or “combined phenotype” according to the site of main dilatation. Baseline clinical and echo data were compared according to aortic valve function (AS-BAV vs BAV with normal valve function, AR-BAV vs BAV with normal valve function, AS-BAV versus AR-BAV).

### Human aortic valve samples and processing

The protocol was evaluated and authorized by the “CPP Sud Méditerranée IV” n°:13.061 and by the “Agence de la biomedicine” n°PFS14-011. Aortic valve leaflets were collected from patients with L-R BAV with raphe operated on for severe AS or AR in our cardiac surgery department. Normal tricuspid aortic valve were collected on human heart explant immediately after transplantation from patients with endstage of heart failure and no aortic valve disease. Immediately following surgical removal, samples were immersed in *RNAlater* solution (Sigma) and stored at −80°C. Tissue was lysed with TRIzol™ reagent (Thermo Fisher Scientific), and total RNAs were purified with the RNeasy mini Kit (Qiagen). Reverse transcriptions were performed using first strand cDNA synthesis kit (Roche) following the manufacturer’s instructions. LightCycler 480 SYBR Green I Master mix (Roche) was used for quantitative real-time PCR (RT-qPCR) analysis with a LightCycler 480 (Roche) following the manufacturer’s instructions. The genetic study focused on several genes involved in valve development and extracellular matrix homeostasis to compare their expression according to valve dysfunction. Among genes with significant difference of expression between AS-BAV and AR-BAV (n= 15) *RUNX2, COL1A1* and *FLNA* were chosen due to their proven role in extracellular valve remodeling and calcification process.

The following primers were used for the RT-qPCR:

**RUNX2 primers:** *Forward: 5’-CAGGCAGGTGCTTCAGAACT-3’; Reverse: 5’-GACTGGCGGGGTGTAAGTAA-3’.*
**COL1A1 primers:** *Forward: 5’-CTGGATGCCATCAAAGTCT-3’; Reverse:5’-TCTTGTCCTTGGGGTTCTTG-3’.*
**FLNA primers:** *Forward: 5’-AAGTGACCGCCAATAACGAC-3’; Reverse: 5’-CCAGCAAAGAGCACAGTAACC-3’.*
**GAPDH primers:** *Forward: 5’-GTCCACTGGCGTCTTCACCA - 3’; Reverse: 5’-GTGGCAGTGATGGCATGGAC - 3’*

RT-qPCR was performed in triplicate for each genotype. Samples were normalized to *GAPDH* as endogenous housekeeping gene. All tests were performed two-sided and for all analyses, p<0.05 was considered statistically significant. For gene expression comparison, mRNA expression levels were calculated using the comparative cycle threshold(ΔΔCT) method. Normalized expression levels in the normal tricuspid aortic valve were set to 1.0 and were considered as control. Expression level in AS-BAV and AR-BAV was first compared to expression in healthy tricuspid valve and then to each other. Statistical analysis was carried out using student *t* test.

### Statistical analysis

Statistical analyses were performed using R-software 3.0.3. Quantitative variables are expressed as mean±SD and categorical variables are presented as numbers and percentages. Comparisons between groups were performed using the Student’s t test or the Mann-Whitney U test for quantitative variables, and using the chi-2 test or the Fisher’s test for categorical variables. Univariate logistic regression analyses were performed to assess the association between potential determinants of valve dysfunction. Firth’s correction was applied by performing Firth’s penalized-likelihood logistic regression to take into account the small number of patients without valve dysfunction.^13^ Potential factor associated with valve dysfunction were considered as candidate for the multivariable analysis when their p-value was less than 0.20 according to the univariate analysis. A backward selection procedure was then applied to build a final regression model.

## RESULTS

### Baseline characteristics (Table 1)

**Table 1:** Baseline clinical characteristics.

|  | Aortic valve function at inclusion |  |  |  |  |  |  |
| --- | --- | --- | --- | --- | --- | --- | --- |
|  | Overall | Normal valve function | AS | AR | AR+AS | p value |  |
|  | n=223 | n=24 | n=68 | n=91 | n=40 | NF vs dysf | AS vs.AR |
| Age,years | 53±17 | 42±15 | 66±10 | 44±14 | 56.2±13 | <0.01 | <0.01 |
| Age>50years(%) | 126(56.5%) | 6(25%) | 62(62.4%) | 30(32.9%) | 29(72.5%) | <0.01 | <0.01 |
| Male gender(%) | 168(75.3%) | 14(58.3%) | 45(60.6%) | 78(86%) | 31(77.5%) | 0.04 | 0.06 |
| BSA(m2) | 1.89±0.23 | 1.82±0.28 | 1.85±0.22 | 1.93±0.24 | 1.9±0.21 | 0.44 | <0.01 |
| Diabetes(%) | 13(5.8%) | 0(0) | 10(14.7%) | 1(1.1%) | 2(5%) | 0.28 | <0.01 |
| Hypertension(%) | 67(30%) | 7(29.2%) | 29(42.6%) | 20(21.9%) | 11(27.5%) | 0.9 | <0.01 |
| Dyslipidemia(%) | 43(19.3%) | 1(4.1%) | 26(38.2%) | 8(8.8%) | 8(20%) | 0.4 | <0.01 |
| Coronary artery disease(%) | 32(14.3%) | 1(4.1%) | 17(25.7%) | 5(5.5%) | 9(22.5%) | 0.29 | <0.01 |
| Renal insufficiency(%) | 1(0.4%) | 0(0) | 0(0) | 1(1.1%) | 0 (0) | 0.09 | 0.86 |
| Chronic pulmonary disease | 4(1.8%) | 0(0) | 1(1.5%) | 2(2.2%) | 1(2.5%) | 0.3 | 0.3 |
| Stroke(%) | 11(4.9%) | 0(0) | 5(5.6%) | 4(4.4%) | 2(5%) | 0.4 | 0.5 |
| Atrial fibrillation(%) | 10(5.8%) | 3 (12.5%) | 3(4.5%) | 3(3.3%) | 1(2.5%) | 0.23 | 0.65 |
| CHF(%) | 29(13%) | 0(0) | 10(15.2%) | 9(9.9%) | 10(25%) | 0.04 | 0.48 |
| Endocarditis(%) | 17(7.6%) | 1 (4.1%) | 1(1.5%) | 11(12.1%) | 3(7.5%) | 0.18 | <0.01 |
| Aortic Coarctation(%) | 8(3.6%) | 1(4.1%) | 0(0) | 5(5.5%) | 2(5%) | 0.31 | <0.01 |
| Aortic Dissection(%) | 0(0%) | 0(0) | 0(0) | 0(0) | 0(0) | 0 | 0 |
| Familial history of BAV(%) | 25(11.2%) | 4(16.6%) | 4(6.6%) | 11(12%) | 6(6.6%) | 0.4 | <0.01 |
| NYHA I/II | 181(81.1%) | 24(100%) | 49(74.2%) | 75(82.4%) | 33(82.5%) | 0.03 | 0.4 |
| BNP(ng/mL) | 151.3±228.7 | 65.2±68.4 | 138.5±155.6 | 147±284.7 | 207±264.5 | <0.01 | 0.6 |

Two hundred twenty three patients were prospectively included in the cohort Mean age was 53±17years, majority were asymptomatic and male (sex ratio = 3:1). None had history of aortic dissection or family history of unexplained sudden death in the youth. Among 17 patients with infective endocarditis (IE) at presentation, 12(5% of all BAV patients) had isolated aortic valve endocarditis and 4 (24% of all IE) peri annular complications. Cardiac surgery was performed within 6 months of diagnosis in 145 patients (65%), including 141 aortic valve replacements, mainly with bio-prothesis(n=85) and 4 isolated ascending aorta tubular graft replacement. Mean age at cardiac surgery was 56±15years and 72(49%) of operated patients were so before the age of sixty. Indication for surgery was AR in 26 patients (18% of all aortic valve replacements, mean age= 44±17 years), AS in 92 patients (65%, mean age=62±13years) and ascending aorta dilatation in 27 patients (including 23 combined surgery).

### BAV phenotype (Table 2)

**Table 2:** Baseline echocardiographic characteristics.

|  | Overall | Normal valve function | Aortic valve function |  |  | p value |  |
| --- | --- | --- | --- | --- | --- | --- | --- |
|  |  |  | AS | AR | AR+AS | NF vs dysf. | AS vs.AR |
|  | n=223 | n=24 | n=68 | n=91 | n=40 |  |  |
| LVEF(%) | 62±10 | 60±8 | 61.7±11 | 62.4±8.2 | 62±11% | 0.71 | 0.85 |
| LV diastolic diameter(mm) | 52±11 | 48±7 | 46.3±8.1 | 54.6±10.1 | 52.8±11.2 | 0.18 | <0.01 |
| LV systolic diameter(mm) | 35±10 | 32±8 | 31.2±10.1 | 37.28.5 | 34.8±10.8 | 0.66 | 0.08 |
| LV diastolic volume(ml) | 161±72 | 131±49 | 116.8±46.7 | 196.3±71.7 | 157 | 0.04 | <0.01 |
| LV systolic volume(ml) | 67±43 | 54±38 | 52.4±33.7 | 80.6±48 | 71.3 | <0.01 | <0.01 |
| Aortic annulus diameter(mm) | 26±4 | 26±3 | 24.7±3.7 | 27.2±4 | 25.2±3.6 | 0.70 | <0.01 |
| Presence of raphe(%) | 182(82%) | 14(54%) | 62(91.2) | 76(83.5) | 30(75%) | <0.01 | 0.32 |
| **Sievers Classification** |  |  |  |  |  | 0.23 | 0.42 |
| Type 0(%) | 34(15.2%) | 6(25%) | 6(8.8%) | 13(14.3%) | 9(22.5%) |  |  |
| Type 1(%) | 178(79.8%) | 13(54.2%) | 60(88.2%) | 75(82.4%) | 30(75%) |  |  |
| Type 1 with R/L Fusion(%) | 157(70.4%) | 12(50%) | 55(80.9%) | 66(72.5%) | 24(60%) |  |  |
| Type 2(%) | 4(1.8%) | 1(4.2%) | 2(3%) | 1(1.1%) | 0 |  |  |
| Unknown(%) | 7(3.2%) | 4(16.6%) | 0 | 2(2.2%) | 1(2.5%) |  |  |
| **Embryologic Classification** |  |  |  |  |  | 0.58 | 0.91 |
| Typical phenotype L/R(%) | 186(83.4%) | 18(75%) | 60(88.2%) | 77(84.6%) | 31(77.5%) |  |  |
| Atypical phenotyppe NC/R(%) | 23(10.3%) | 1(4.1%) | 5(7.3%) | 10(11%) | 7(17.5%) |  |  |
| Atypical phenotyppe NC/L(%) | 3(1.3%) | 0(0) | 1(1.5%) | 1(1.1%) | 1(2.5%) |  |  |
| Unknown(%) | 11(5%) | 5(20.9%) | 2(3%) | 3(3.3%) | 1(2.5%) |  |  |
| **BAV aortopathy (%)** | **110(49.3%)** | **12(50%)** | **21(30.8%)** | **55(60.4%)** | **22(55%)** | 0.41 | <0.01 |
| Aortic root diameter(mm) | 40±8 | 43±18 | 37.2±5.8 | 32.5±5.8 | 38.2±6.7 | 0.42 | 0.63 |
| Ascending aorta diameter(mm) | 42±9 | 44±16 | 41.5±6.8 | 41.3±7.6 | 41.7±8.3 | 0.69 | 0.35 |
| Root phenotype(%) | 49/110(44.5%) | 6/12(50%) | 1/21(5%) | 39/55(71%) | 3/22(13.6%) | 0.6 | <0.01 |
| Tubular phenotype(%) | 51/110(46.5%) | 4(33.3%) | 19/21(90%) | 16/55(29%) | 12/22(54.6%) | 0.38 | <0.01 |
| Combined phenotype(%) | 10/110(9%) | 2(16.7%) | 1/21(5%) | 0(0) | 7/22(31.8%) | 0.69 | 0.9 |

According to Sievers classification, majority of patients (80%) had type 1 BAV, localized between left and right coronary cusps in 88% of them, 15% had type 0 and only 2% had type 2 BAV. According to embryologic classification, L-R BAV with or without raphe was observed in 83% of all BAV patients, NC-R BAV in 10% and NC-L BAV in 1%. Aortopathy was present at diagnosis in half BAV patients, evenly distributed between root and tubular phenotype.

### Normal valve function (Tables 1 and 2)

Twenty-four patients (11% of all patients) had normal valve function and 199 patients (89%) presented with valvular dysfunction at the time of inclusion. Patients with normal valve function were predominantly male, younger than patients with AS or combined AS+AR (p<0.01) but were of similar age than patients with isolated AR at diagnosis (p=0.4). Only four of those patients with normal valve function were older than 60 years old. They presented less frequently with raphe than patients with valve dysfunction (54%vs.84%,p<0.01). Majority had L-R BAV as patients with valve dysfunction (75%vs.84%,p=0.58). Prevalence of aortopathy was similar to that observed in patients with any type of valve dysfunction (50%vs.49%,p=0.41).

### Aortic stenosis (Tables 1 and 2)

Sixty-eight patients (31% of all patients) had isolated AS, including 58(26% of all BAV patients) with severe AS. Mean indexed aortic valve area was 0.49±0.24 cm^2^/m^2^ and mean trans-valvular gradient was 56±21mmHg. Patients with AS were significantly older than patients with normal function or with isolated AR-BAV. They were predominantly male but proportion of women (39%) was higher than among AR patients (14%). Majority had raphe and L-R BAV as patients with AR. Prevalence of aortopathy was lower than among patients with isolated AR and of different phenotype, mostly tubular.

### Aortic regurgitation (Tables 1 and 2)

Ninety-one patients (41% of all patients) had isolated AR, including 50 (22% of all BAV patients) with severe AR. Mean effective regurgitant orifice area 34±15mm^2^ and mean regurgitant volume was 83±39ml. As expected, LV diameters and volumes were higher than in AS-BAV patients. Mechanism of AR was pure leaflet prolapse in 40 patients (44% of all AR patients), restriction in 5 (6%), combined prolapse and restriction in 46 (50%). Among AR-BAV patients, male predominance was more prominent than among AS-BAV patients, mean age was 22 years younger, comorbidity was lower but endocarditis higher at diagnosis. Majority had raphe and L-R BAV. Prevalence of aortopathy was twice higher than among patients with AS, mostly of root phenotype.

### Combined AS and AR

Forty patients (18%) had combined significant AS and AR. They were predominantly male and clinical and echocardiographic pattern was in-between isolated AS and AR patients. Majority had raphe and L-R BAV. Aortopathy was mostly of tubular or combined phenotype.

### Comparison between AS-BAV and BAV with normal valve function

By univariate analysis (OR[95%CI]), age>50 years (18.6[6.2-64],p<0.001), presence of raphe (5.7[1.8-19.5], p=0.003), diabetes (8.8[1.1-1147.3],p=0.04) and dyslipidemia (3.8[1.2-15.5],p=0.02) were significantly associated with BAV stenosis whereas male gender (1.4[0.5-3,6],p=0.5) and orientation of commissures (1.5[0.3-14.8],p=0.7) were not. By multivariable analysis, age>50 years (41.6[10.3-248.2],p<0.001) and presence of a raphe (12.8[2.4-87.4],p<0.001) were significantly associated with AS.

### Comparison between AR-BAV and BAV with normal valve function

By univariate analysis, male gender (5.0[1.6-16.4],p=0.005), aortic annulus diameter(1.2[1.02-1.56],p=0.03) and presence of raphe (3.6[1.1-11.9],p=0.03) were significantly associated with BAV regurgitation whereas age (0.9[0.3-2.5],p=0.8) and orientation of commissures (1.9[0.4-19.0],p=0.5) were not. By multivariable analysis, only male gender (5[1.6-16.4],p=0.005) was significantly associated with AR.

### Comparison between AR-BAV and AS-BAV

By univariate analysis, male gender (3.6[1.7-8.7],p=0.001), younger age (10[5.2-20.0],p<0.001) and higher aortic annulus diameter (1.27[1.14-1.43],p=0.001) were significantly associated with AR whereas dyslipidaemia (3.3[1.6-7.1,p=0.002) was associated with AS. Multivariable analysis identified younger age (1.9[1.85-1.94],p<0.001) as independently associated with AR and dyslipidemia (3.3[1.22-8.22],p<0.001) with AS, while male gender was of borderline significance for predicting AR (1.15 [0.70-2.54],p=0.07).

### Comparison of gene expression level in AS-BAV and AR-BAV

To compare expression levels in AS-BAV and AR-BAV we performed RT-qPCR on surgical specimens with AS-BAV (n=7) and AR-BAV (n=8). Healthy tricuspid aortic valves collected on human heart explant immediately after transplantation were used as controls (n=5). A total of 20 aortic valve samples were analyzed.

All BAV patients had L-R BAV with raphe. AS-BAV patients included 5 males, mean age was 72±5 and surgical correction was indicated for severe symptomatic AS. Mean gradient was 57±14mmHg, mean LVEF was 65±10%. Aortopathy was present in 4 patients, of tubular phenotype in 2. AR-BAV patients included 8 males, mean age was 40±13 and surgical correction was indicated for severe symptomatic AR. Mean ERO was 35±11mm^2^, mean regurgitant volume was 70±13ml and mean LVEF=62±12%. Aortopathy was present in 6 patients, of root phenotype in all.

Tricuspid aortic valves used as controls were collected from five patients with end-stage congestive heart failure due to dilated or ischemic cardiomyopathy. Male gender was prevalent (n=3, 60%) and mean age was 53±5.

Transcription level of *RUNX2*, a downstream transcriptional regulator of osteoblast cell fate,^2^ was significantly increased in AS-BAV compared to controls and to AR-BAV (17.67±1.83vs 3.25±0.93,p=0.04)(**Figure 1**). Transcription level of *COL1A1*, a fibrillar collagen gene expressed in valvular extracellular matrix,^14^ was increased in both AR and AS-BAV patients as compared to controls, but overexpression was significantly more prominent in AR-BAV patients (4.01±0.6 vs 2.25±0.5,p=0.03). Lastly, transcription level of *FLNA*, which encodes a cytoskeletal protein that has been implicated as critical mediator of cell signaling and migration^15^, was significantly increased in AR-BAV patients as compared to controls and to AS-BAV patients (23.31±7.5 vs 1.97±0.3,p=0.03).

**Figure:**
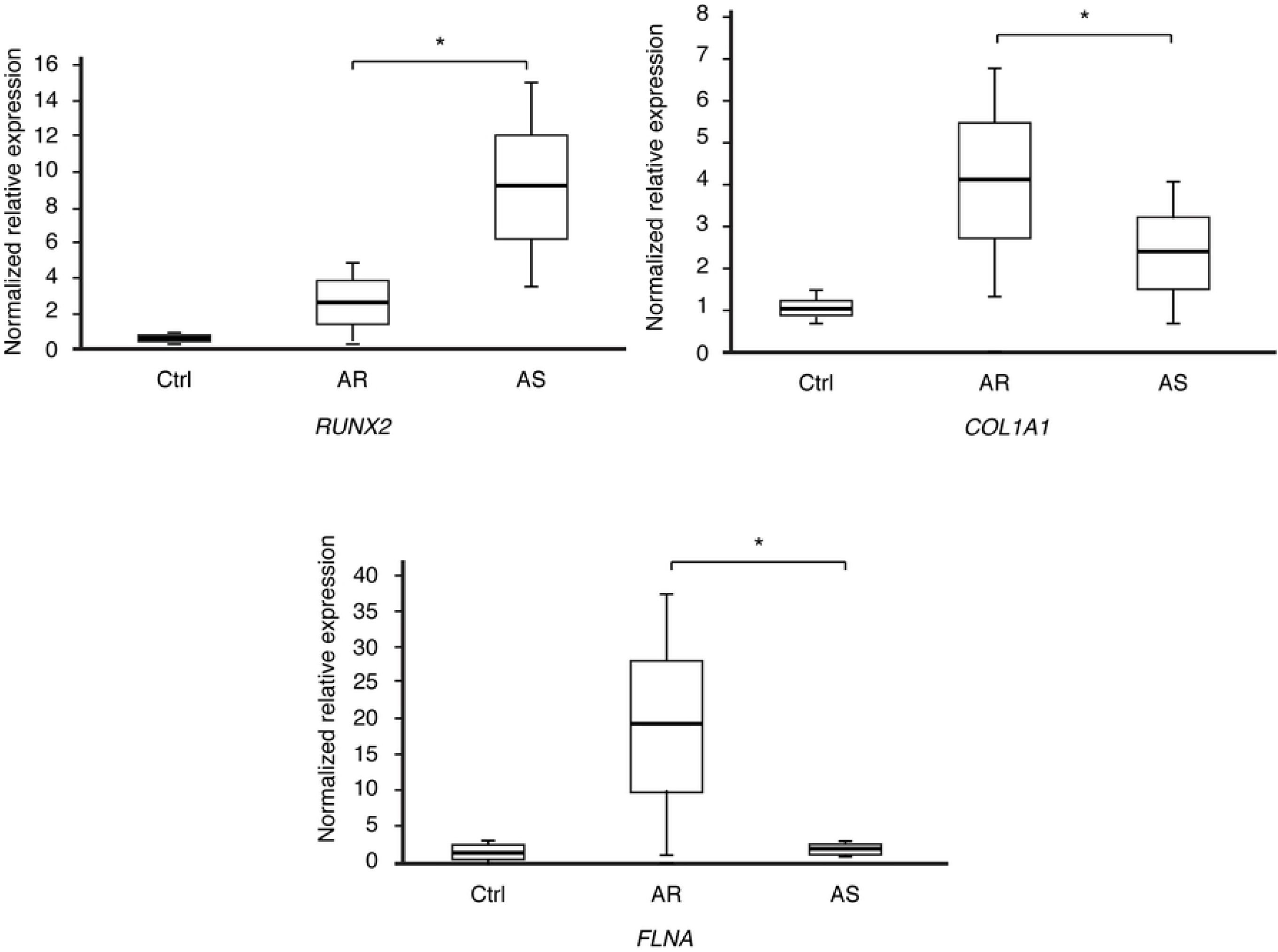
Comparison of valve tissue transcription levels of *RUNX2, COL1A1* and *FLNA* between normal aortic valve controls, AR-BAV and AS-BAV patients.

## DISCUSSION

The present study conducted on 223 consecutive patients diagnosed with BAV in routine clinical practice shows that in a tertiary care center 1) prevalence of severe AS and AR are similar, observed in a quarter of all patients at diagnosis 2) AR is mainly due to leaflet prolapse and occurs at a much younger age than AS 3) presence of raphe is associated with valve dysfunction mainly AS whereas embryologic phenotype is not 4) the leaking or stenotic processes could be both supported by deregulation of specific genetic pathways

BAV is the most frequent CHD and is associated with high rates of valve degeneration.^7,8,16^ Development of ascending aorta aneurysm is observed in one fourth of patients 25 years after diagnosis of BAV with an excess risk of aortic dissection.^8^ Subsequently BAV patients experience high rates of surgical aortic interventions at younger age than patients with tricuspid aortic valve.^17,7,6,8^ However, data from populations referred to tertiary care centers at evolving clinical and echocardiographic stages unveil excess mortality in some subsets as compared to expected survival in an age- and sex-matched population due to higher prevalence of valvular dysfunction.^3^ Our study is consistent with these data^3^ reporting low prevalence of strictly normally functioning aortic valve at diagnosis of BAV, barely exceeding 10% of patients despite the young age of the study population and the exclusion of patients directly referred for surgery. Almost none of them were older than 60, making valve degeneration an early process. Conversely, the vast majority of BAV patients presented with significant valvular dysfunction at diagnosis, associated with aortopathy in half of them, explaining the high incidence of surgery within 6 months of diagnosis reaching two thirds of patients at a mean age of 56 years-old, even younger when indication was driven by AR.

Such high burden of morbidity and mortality imposes comprehensive assessment of determinants of clinical outcome and mechanisms of valve degeneration, critical for follow-up and management of patients with a first diagnosis of BAV. To date, identified risk factors of cardiac events, include presence of valve degeneration at first diagnosis of BAV, baseline ascending aorta diameter>40mm,^7^ female gender particularly with significant AR^3^ and older age.^3,7^ In turn, risk factors of outcome in patients with strictly normal BAV are poorly defined and BAV ability to maintain normal valve function or to become dysfunctional is not understood. BAV phenotype has been correlated to the type of dysfunction in one study, typical phenotype, defined by 11-17 or 10-16 o’clock positions of commissural axis, being associated with AR and atypical phenotype, defined by 13-19 or 12-18 o’clock positions of commissural axis,being associated with AS.^18^ Our results do not confirm this association since embryologic phenotype was not associated with neither type of dysfunction, which if consistent with the overwhelming majority of L-R fusion in community and tertiary series^3^, making hazardous any attempt of risk stratification using this parameter. Proportions of typical and atypical phenotype were indeed almost identical in our series in both types of degeneration and close to those observed in normal BAV. Likewise, within each phenotype, normal valve function, leaking or stenotic degeneration were evenly distributed. Presence of raphe has been associated with higher prevalence of AS and AR-BAV and increased incidence of aortic surgery.^19^ BAV differs from the tricuspid valve by number of cusps and sinus, by the elliptical shaped of the aortic ring and the restrictive and asymmetrical opening of aortic valve during systole.^20,21^ MRI revealed eccentric systolic jet and abnormal downstream helical flow patterns through normal BAV resulting in greater flow acceleration and higher shear stress.^12,22^ This phenomenon attributed to commissural fusion could lead to higher sensitivity to valve degeneration and calcification^12^ and could be magnified by the increased stiffness of the fused leaflet due to the presence of raphe. Our data agree with this hypothesis by showing an independent association between raphe and valve dysfunction mainly AS and a lower prevalence of raphe in normal BAV. However, raphe can be seen as the expression of an already ongoing degenerative process and could be acquired.^23^ In addition, in our population more than half of normally functioning BAV presented with raphe and almost 20% of AR patients without, leaving unanswered the issue of valve dysfunction in many patients and imposing to identify determinants of degeneration independently of phenotype.

The most striking observation of our series was the major age difference at diagnosis of AR as compared to AS-BAV. Patients with significant AR were more than 20 years younger than patients with significant AS, suggesting that both degenerative processes could follow specific genetic pathways. Within the most common BAV phenotype, i.e. L-R BAV with raphe, we observed substantial differences in tissue transcription levels of three signaling pathways according to type of valve dysfunction as compared to normal aortic valve.^24^ Overexpression of *RUNX2* signal observed in our AS BAV patients is consistent with its role on osteogenesis and with its upregulation observed in mice models of calcified aortic valve.^25^ This transcription factor is regulated by NOTCH1, a single-pass transmembrane receptor expressed in valvular endothelial cells during development and involved in prevention of valve calcification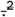 In our AR-BAV samples, the gene expression profile was significantly different with considerably lower overexpression of *RUNX2* and an upregulation of *COL1A1* and *FLNA* genes. *COL1A1* along with *COL1A2*, which are the major components of type I collagen in the aortic leaflets are responsible for the stiffness of the valve.^26^ The upregulation of *COL1A1* is consistent with the reorganization of the extracellular matrix of the aortic leaflets. *FLNA* is the main isoform of filamins, which are proteins stabilizing actin cytoskeleton playing a crucial role in the interactions between valvular interstitial cells and their environment. Mutations in *FLNA* gene have been linked to mitral valve prolapse.^27,28^ Altogether, dysregulation of these last two genes could be responsible for remodeling and disruption of the extracellular matrix, embarking aortic leaflets towards a leaking process which histologic basis could be close to myxomatous degeneration. This is in line with leaflet prolapse being the main mechanism for AR in our BAV patients. Expression of *RUNX2* gene in AR patients is compared to controls. Lastly, location of *FLNA* on the X chromosome could be coherent with the higher male predominance observed among our AR-BAV patients.

### Limitations

Any cohort of BAV patients is susceptible to diagnostic errors between tricuspid and bicuspid valve, with the risk of data contamination by non BAV patients. The prospective design of this study and the cautiousness taken in scanning with exclusion of all unclear diagnosis particularly among AS patients should reduce this bias. The fact that many patients with well-functioning BAV were not diagnosed and not captured in this tertiary cohort creates also a selection bias. Close follow-up of BAV patients from birth to occurrence of valve dysfunction would be the ideal way to disclose mechanisms of degeneration but is not conceivable. Therefore, cross sectional comparison of BAV patients according to valve function at first diagnosis of BAV is an imperfect but acceptable method.

Our genetic study was performed on a small number of tissue samples due to the low number of samples that have passed our quality control for RNA extraction. However, this cautiousness a warrant of the homogeneity of our results.

Valvular degeneration is obviously a complex interplay of various factors including hemodynamic constraints due to the fused leaflets and genetic pathways dysregulation, leading to maladaptive response of valvular extracellular matrix to the aortic flow environment. It is unknown whether up or down regulations of signaling pathways are genetically programmed and in connection with the genetic background of BAV itself or triggered by postnatal environmental factors. Both hypotheses are plausible since NOTCH1 mutations have been reported to explain both the occurrence of BAV and its early degeneration.^2^ Similarly, age marked differences at initiation of leaking or stenotic process are unclear. Specific interactions between genetic pathways and environmental factors linked to aging could be argued. Finally, myxomatous degeneration as the histologic basis of AR BAV is attractive but remains to be definitely proven by larger data.

## CONCLUSION

This study confirmed high prevalence of valve dysfunction at first diagnosis of BAV among patients referred in a tertiary care center. The leaking or stenotic processes are poorly associated with clinical and phenotypic markers. This emphasize that BAV degeneration is a complex phenomenon which involves several contributors such as environmental factors shared with tricuspid aortic valve and local hemodynamic factors specific to BAV. Dysregulation of specific genetic pathways could play a role and need to be thoroughly investigated.

## REFERENCES

1. Sievers H-H, Schmidtke C. A classification system for the bicuspid aortic valve from 304 surgical specimens. J Thorac Cardiovasc Surg. 2007;133:1226–1233.

2. Garg V, Muth AN, Ransom JF, et al. Mutations in NOTCH1 cause aortic valve disease. Nature. 2005;437:270–4.

3. Michelena HI, Suri RM, Katan O, et al. Sex Differences and Survival in Adults With Bicuspid Aortic Valves: Verification in 3 Contemporary Echocardiographic Cohorts. J Am Heart Assoc. 2016;5:e004211.

4. Hoffman JIE, Kaplan S. The incidence of congenital heart disease. J Am Coll Cardiol. 2002;39:1890–900.

5. Larson EW, Edwards WD. Risk factors for aortic dissection: a necropsy study of 161 cases. Am J Cardiol. 1984;53:849–55.

6. Michelena HI, Prakash SK, Della Corte A, et al. Bicuspid aortic valve: identifying knowledge gaps and rising to the challenge from the International Bicuspid Aortic Valve Consortium (BAVCon). Circulation. 2014;129:2691–704.

7. Michelena HI, Desjardins VA, Avierinos J-F, et al. Natural history of asymptomatic patients with normally functioning or minimally dysfunctional bicuspid aortic valve in the community. Circulation. 2008;117:2776–84.

8. Michelena HI, Khanna AD, Mahoney D, et al. Incidence of aortic complications in patients with bicuspid aortic valves. JAMA. 2011;306:1104–12.

9. Baumgartner H, Falk V, Bax JJ, et al. ESC Scientific Document Group. 2017 ESC/EACTS Guidelines for the management of valvular heart disease. Eur Heart J. 2017;38:2739–2791.

10. Nishimura RA, Otto CM, Bonow RO, et al. 2017 AHA/ACC Focused Update of the 2014 AHA/ACC Guideline for the Management of Patients With Valvular Heart Disease. J Am Coll Cardiol. 2017;70:252–289.

11. Atkins SK, Sucosky P. Etiology of bicuspid aortic valve disease: Focus on hemodynamics. World J Cardiol. 2014;6:1227–33.

12. Sun L, Chandra S, Sucosky P. Ex vivo evidence for the contribution of hemodynamic shear stress abnormalities to the early pathogenesis of calcific bicuspid aortic valve disease. PLoS One. 2012;7:e48843.

13. Firth D. Bias Reduction of Maximum Likelihood. Biometrika. 1993;80:27–38.

14. Odelin G, Faure E, Kober F, et al. Loss of Krox20 results in aortic valve regurgitation and impaired transcriptional activation of fibrillar collagen genes. Cardiovasc Res. 2014;104:443–55.

15. Zhou A-X, Hartwig JH, Akyürek LM. Filamins in cell signaling, transcription and organ development. Trends Cell Biol. 2010;20:113–123.

16. Michelena HI, Mankad S V. Sex Differences in Bicuspid Aortic Valve Adults: Who Deserves Our Attention, Men or Women? Circ Cardiovasc Imaging. 2017;10:e006123.

17. Roberts WC, Ko JM. Frequency by Decades of Unicuspid, Bicuspid, and Tricuspid Aortic Valves in Adults Having Isolated Aortic Valve Replacement for Aortic Stenosis, With or Without Associated Aortic Regurgitation. Circulation. 2005;111:920–925.

18. Kang J-W, Song HG, Yang DH, et al. Association Between Bicuspid Aortic Valve Phenotype and Patterns of Valvular Dysfunction and Bicuspid Aortopathy. JACC Cardiovasc Imaging. 2013;6:150–161.

19. Kong WKF, Delgado V, Poh KK, et al. Prognostic implications of raphe in bicuspid aortic valve anatomy. JAMA Cardiol. 2017;2(3):285–292

20. Bäck M, Gasser TC, Michel J-B, Caligiuri G. Biomechanical factors in the biology of aortic wall and aortic valve diseases. Cardiovasc Res. 2013;99:232–41.

21. Robicsek F, Thubrikar MJ, Cook JW, Fowler B. The congenitally bicuspid aortic valve: how does it function? Why does it fail? Ann Thorac Surg. 2004;77:177–185.

22. Marom G, Kim H-S, Rosenfeld M, Raanani E, Haj-Ali R. Fully coupled fluid-structure interaction model of congenital bicuspid aortic valves: effect of asymmetry on hemodynamics. Med Biol Eng Comput. 2013;51:839–48.

23. Yang L-T, Enriquez-Sarano M, Michelena HI. The bicuspid aortic valve raphe: an evolving structure. Eur Hear J - Cardiovasc Imaging. 2020;21:590–590.

24. Rajamannan NM. CALCIFIC AORTIC STENOSIS: FROM BENCH TO THE BEDSIDE-EMERGING CLINICAL AND CELLULAR CONCEPTS. Heart. 2003;89:801–805.

25. Steitz SA, Speer MY, Curinga G, et al. Smooth muscle cell phenotypic transition associated with calcification: Upregulation of Cbfa1 and downregulation of smooth muscle lineage markers. Circ Res. 2001;89(12):1147–54

26. Leopold JA. Cellular mechanisms of aortic valve calcification. Circ Cardiovasc Interv. 2012;5:605–14.

27. Trochu J-N, Kyndt F, Schott J-J, et al. Clinical characteristics of a familial inherited myxomatous valvular dystrophy mapped to Xq28. J Am Coll Cardiol. 2000;35:1890–1897.

28. Kyndt F, Gueffet J-P, Probst V, et al. Mutations in the Gene Encoding Filamin A as a Cause for Familial Cardiac Valvular Dystrophy. Circulation. 2007;115:40–49.

